# Age-related decline in function of ON and OFF visual pathways

**DOI:** 10.1101/2021.12.03.471168

**Authors:** Amithavikram R Hathibelagal, Vishal Prajapati, Indrani Jayagopi, Subhadra Jalali, Shonraj Ballae Ganeshrao

## Abstract

**Purpose:** Simple psychophysical paradigm is available as a digital application in iOS devices such as iPad to measure the function of ON and OFF visual pathways. However, an age-matched normative database is not readily available. The purpose of the study is to evaluate the response of ON and OFF visual pathways as a function of age.

**Methods:** 158 normal healthy adults (84 males and 74 females) whose age ranged 18-80 years participated in the study. None of them had any ocular disease (except cataract of grade II or less) and visual acuity of ≤ 20/25. Monocular testing (only one eye) was performed on the ‘EyeSpeed’ application on an iPad at 40cm distance. The targets ranged between 1 to 3 light or dark squares presented randomly in a noise background and participants responded by indicating the number of squares by touching the screen as fast as possible. The main outcome variables are reaction time, accuracy and performance index (1 / speed * accuracy).

**Results:** The median reaction time was shorter (Median (IQR): 1.53s (0.49) [dark] Vs 1.76s (0.58) [light], p < 0.001) and accuracy was higher (97.21% (3.30) [dark] Vs 95.15% (5.10) [light], p < 0.001) for dark targets than the light targets. Performance index and reaction time for both target types significantly correlated with age (ρ = −0.41 to −0.43; p < 0.001).

**Conclusions:** This normative database will be useful to quantify disease-specific defects. More importantly, the ON pathway function can potentially serve as a surrogate for rod photoreceptor function.

## Introduction

The luminance information from the eye to the brain is carried out by two pathways namely ON and OFF pathways [1], which starts in the outer retina at the level of bipolar cells [2]. The ON pathway is responsible for the transmission of signals at/during light stimulation, whereas the OFF pathway is responsible for the transmission of signals after the light offset [1]. The neuronal and cortical resources allocated towards the OFF pathway is more than the ON pathways [3–5] which probably results in more accurate and faster detection of dark targets compared to light targets [6–8]. Beyond animal studies,[3–5] this ON-OFF asymmetry has been demonstrated in humans using simple psychophysical paradigms [6–8]. More importantly, ON/OFF pathway-specific defects have been noted in congenital stationary night blindness [9], melanoma-associated retinopathy [9], amblyopia [10] and glaucoma [8]. However, before using this test as a metric to identify disease-specific defects, one must have an age-matched normative database to distinguish between age-related decline compared to disease-specific defects.

Previously, the effect of age on the function of ON and OFF pathways has been studied in a small subset of individuals (n = 21) between 49-74 years [8]. However, such a small sample is not wide enough to capture the age-related changes due to the following reasons. 1. The age at which visual function begins to decline cannot be captured. 2. The rate of change could be different in different phases of ageing. Besides, the test used previously was lab-based and the testing distance and size of the stimuli are different from the setup used in this study. However, a similar paradigm is now available for testing in a simple portable device as an application. The advantage of having a rapid, sensitive test that is portable is an attractive option in a clinical population. In this study, we aimed to determine the age-related changes in the detection of light and dark targets using an iPad-based freely available application called “EyeSpeed”.

## Methods

This is a prospective cross-sectional study. The study protocol has been approved by the Institutional Review Board (IRB) of the L V Prasad Eye Institute, Hyderabad (IRB Application number: LEC 09-18-139). The inclusion criteria for the participants were as follows: 1. Age ≥ 18 years 2. No history of ocular/systemic disease. 3. Visual acuity at least 20/25 in the tested eye with refractive correction in place. 4. Cataract grade less than LOCS grade II or lower [11]. 5. No history of ocular surgery except pterygium and cataract. All the potential participants were briefed about the test and only if they agreed to participate, then they sign the consent form before participating in the study. The participant cohort included the patient population that visited the hospital campus, also the staff and student cohort from the institute.

### Procedure

The participants wore their near correction if they were ≥ 40 years. The participant’s dominant eye was tested, while the other eye was occluded with a dark patch. The room illumination was standard room illumination of ∼300 lux. The app called “EyeSpeed” was developed by a research group in USA [8] and is available on the iOS app store for free. The application was developed as a tool for easy testing of the ON and OFF visual pathways.

The test was carried out on an iPad (iPad 4^th^ Generation; Model: MD525X/A, USA). The resolution of the device was 2048×1536 pixels. The participants held the tablet with the support of a table. The tablet was held such that there were no major reflections while viewing the screen during the test. The participant’s task was to correctly identify the number of targets against the noise background as fast as possible by tapping on the corresponding button on the touch-sensitive screen (see Fig 1). The participant was initially habituated to a ‘demo’ test for 20 trials before starting the test to avoid any learning effects. The noise squares width was 10 pixels, while the size of the target was 70 pixels. The tablet’s viewing area subtended 27.80°X 20.82° at a test distance of 40cm. The panel of noise background subtended ∼18°at the distance of 40cm, while the target (dark/light) embedded in the noise background subtended ∼2°at the same distance. The ratio of dark area to total area was 50% and the total number of trials was 200. A photometer (Konica Minolta LS-110, Japan) was used to measure the luminance of the white and black targets. The mean luminance of the white patch was 128 ± 0.75 Cd/m^2^, while the luminance of the black patch was 0.19 Cd/m^2^. The mean luminance of the noise background was 63.28 ± 6.16 Cd/m^2^. During the test, the participants viewed either a dark or a light target and the number of targets ranged from 1 to 3 (see Fig 1). In each presentation, the number of targets and type of targets (dark/light) also varied randomly. This type of stimulus has been used previously to elicit the ON and OFF responses in humans.[6,8] More specifically, light targets stimulate the ON pathway, while dark targets stimulate the OFF pathway. There was no fixed duration of the targets, however, they disappeared after the response was obtained from the participant. The average testing time was approximately 4 - 5 minutes. The key outcome variables obtained from the test was accuracy (%) and reaction time (s).

**Figure 1.**
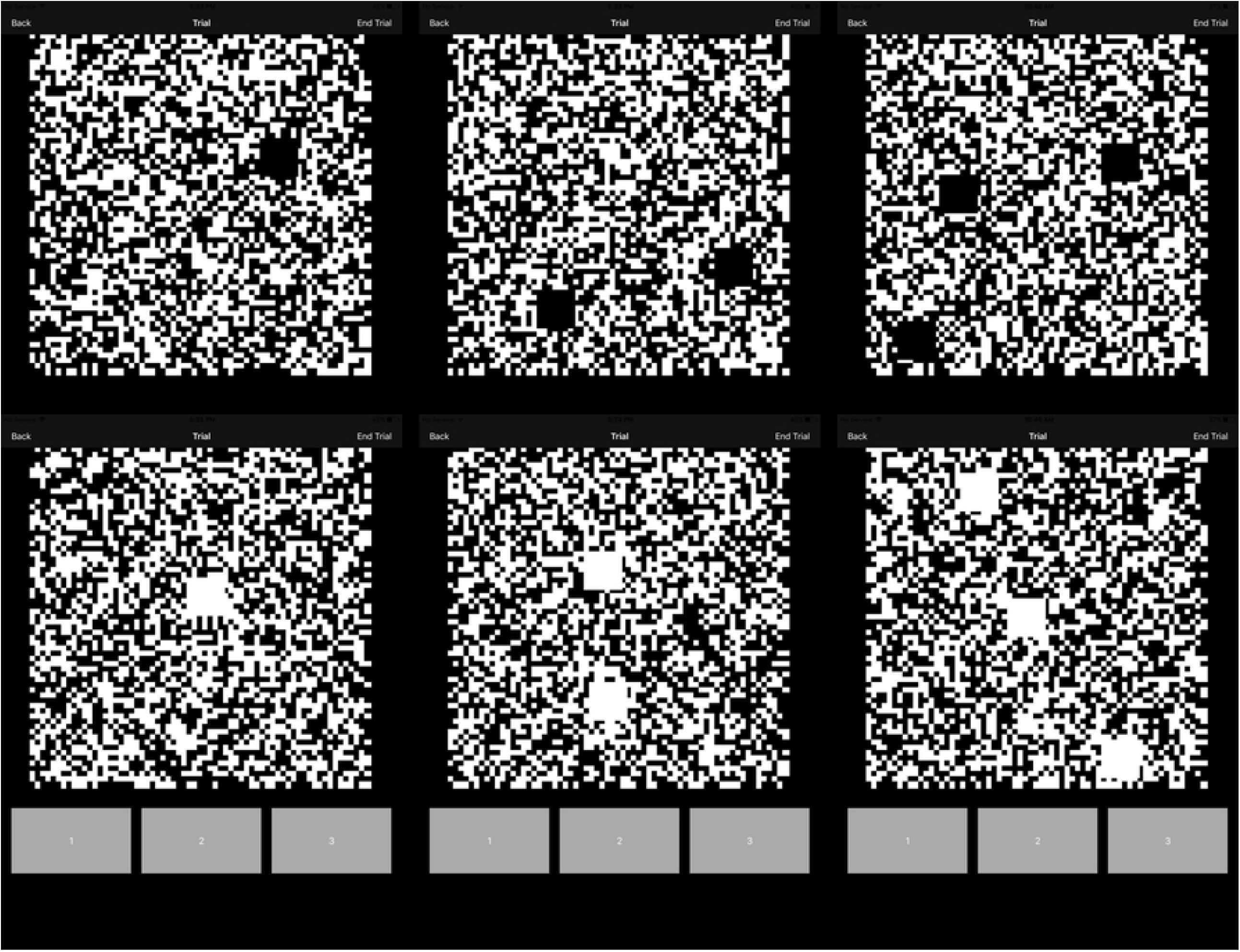
The display screen of the “EyeSpeed” app during the test. The top panels show dark targets in a noise background and the bottom panels show the light targets. The option to choose the correct answer is shown in grey boxes. The number choices are presented in ascending order. Grey option boxes were also present for the top panels as well during the test, which has been removed in this representation for compact viewing.

### Data analysis

The data were not normally distributed as tested by the Kolmogorov-Smirnov test (p > 0.05). Wilcoxon Signed-Rank test is used to compare the outcome parameters between light and dark targets. The p-value of < 0.05 was considered statistically significant. The performance index was computed as the product of speed (1/ reaction time [s]) and accuracy. Linear regression was performed to assess the relationship between age and outcome parameters. The data was divided into 2 age groups namely ≤ 49 years and > 49 years to allow comparison with the previous study [8]. The Grubbs’ test was used to identify the outliers (Graph Pad Prism; https://www.graphpad.com/quickcalcs/Grubbs1.cfm).

## Results

The mean [SD] age of the participants was 43.42 [14.5] years. The repeatability was assessed in a small subgroup of subjects (n = 10). The test was repeated within 1 week from the first visit. The coefficient of variation between visits for all the parameters in both the test conditions was within 20 % of the mean. A total of 170 subjects were approached to participate in the study. Ten subjects (all above 40 years of age) were excluded as there were unable to perform the ‘demo’ test successfully. Two outliers were detected by Grubb’s test (P-value set to 0.05) and were excluded from the analysis. Therefore, data from 158 subjects (84 males and 74 females) were analyzed. The age distribution of subjects is provided in Fig 2. The mean (SD) refractive error of the cohort was −0.36 D (1.29). There were 17 pseudophakes in the cohort (mean (SD) age = 60.29(11.42) years).

**Figure 2.**
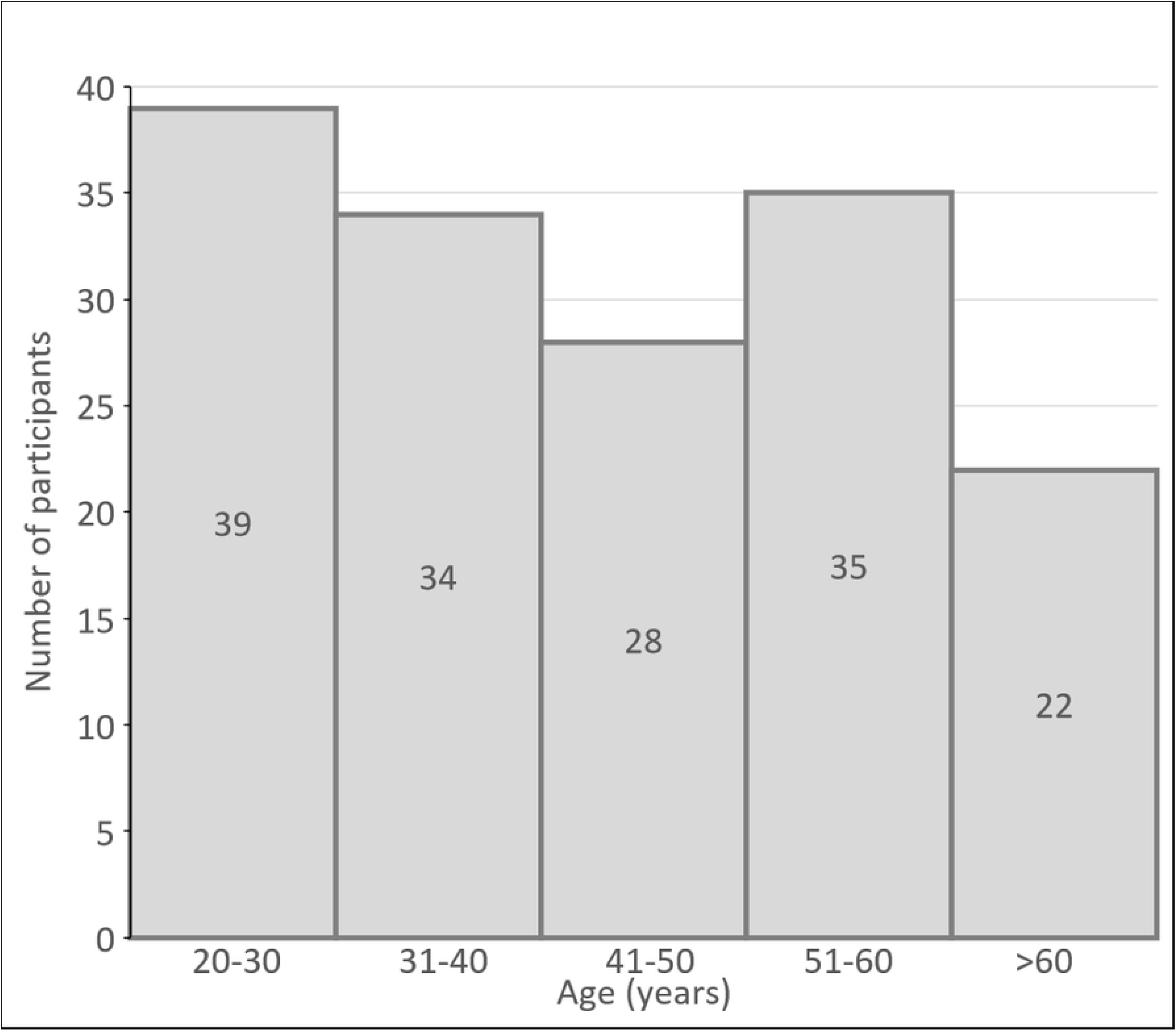
Histogram showing the age distribution of individuals who participated in the study. The numbers inside the bars represent the number of individuals in each decade, with the only exception being the last bar which shows individuals greater than 60 years of age.

The reaction times were shorter (Median (IQR): 1.54s (0.50) [dark] Vs 1.77s (0.63) [light], Wilcoxon Signed Rank test, p < 0.001) and accuracy was higher (97.16% (3.35) [dark] Vs 95.15% (5.32) [light], p < 0.001) for detection of dark targets compared to light targets. Similarly, performance index was also significantly higher for dark targets (63.04%*s^- 1^(20.69)) compared to light targets [(54.3%*s^-1^ (18.27)); p < 0.001]; see Fig 3.

**Figure 3.**
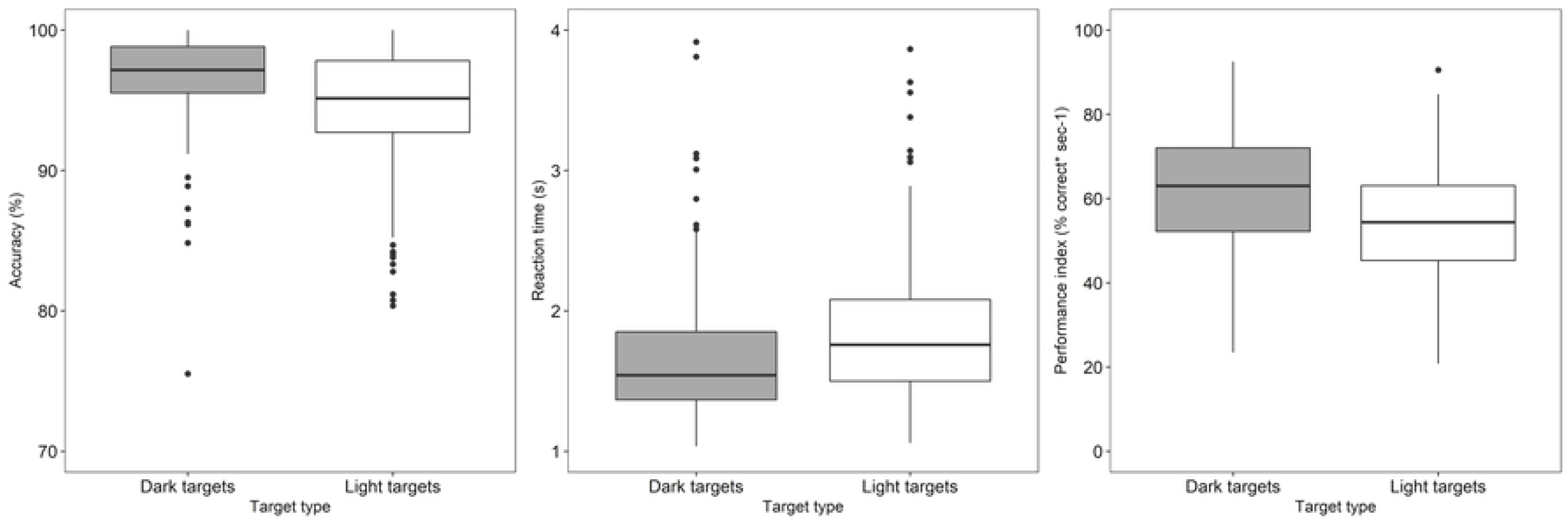
Box plots showing the accuracy (left panel), reaction time (middle panel) and performance index (right most panel) of all the subjects for dark and light targets respectively. The central solid horizontal line in each box represents the median and the solid black circles represent the outliers.

**Figure 4.**
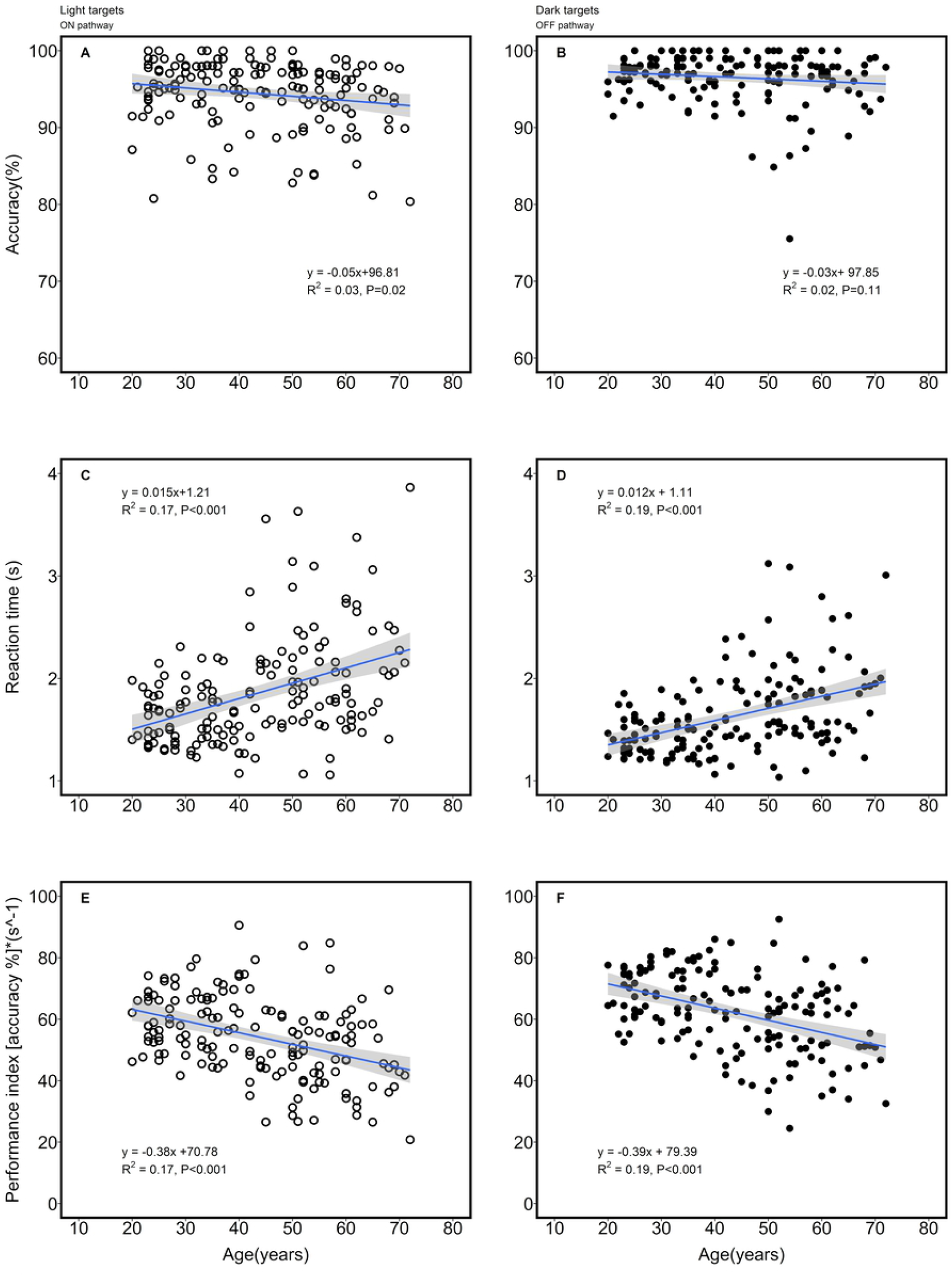
Left panels show outcome variables (accuracy, reaction time and performance index) for light targets as a function of age. A similar representation shows the data for the right panels. The blue solid line in each of the panels represents the slope of the linear regression and the grey region represents the standard error of the fit.

There was no significant difference in accuracy between the two groups age ≤ 49 years and age > 49 years (Table 1). However, there is a significant difference between the reaction times between the two groups, which also manifests as a significant difference in performance index between the two groups. All the outcome parameters showed significant correlation with age except accuracy for black targets (Spearman correlation, ρ = −0.06; p = 0.44) across the entire age range (see Fig 3.). Similarly, the reaction time and performance index measures (for light and dark targets) were significantly correlated (ρ = −0.41 to −0.43; p < 0.001) with age, whereas accuracy (for white targets) showed weak correlation with age (ρ = −0.18; p = 0.02). However, in each of the subgroups (≤ 49 years and age > 49 years), the outcome parameters showed no significant correlation with age. None of the differences in outcome parameters between dark and light targets (e.g., reaction time of dark targets – reaction time of light targets) and age showed any significant correlation. All the outcome parameters were independent of refractive error (p > 0.05). None of the outcome parameters in the IOL group was significantly different from the age-matched cohort. Besides, there were no gender-specific differences in any of the outcome parameters.

**Table 1.** Summary (median [IQR]) of the outcome variables between the two groups

| Outcome parameters | Targets | Age ≤ 49 years (n = 93) | Age > 49 years (n = 65) | p value<br>Mann-Whitney test |
| --- | --- | --- | --- | --- |
| Accuracy (%) | Light | 95.74 [4.19] | 94.59 [6.45] | 0.004 |
|  | Dark | 97.27 [2.95] | 97.06 [3.40] | 0.15 |
| Reaction time (s) | Light | 1.63 [0.48] | 1.97 [0.78] | *0.001 |
|  | Dark | 1.44 [0.33] | 1.80 [0.50] | *0.001 |
| Performance index<br>(accuracy*s<sup>-1</sup>) | Light | 58.47 [17.65] | 46.94 [17.37] | *0.001 |
|  | Dark | 67.88 [15.59] | 54.11 [17.85] | *0.001 |

## Discussion

The main findings from the study are that visual performance in the detection of light and dark targets is dependent on age. Accuracy is independent of age, while the reaction time is relatively constant up to 40 years of age. Beyond this age, there is an increase in reaction time with the increase in age. The higher performance (i.e., shorter reaction times and higher accuracy) for dark targets relative to light targets is consistent with the previous literature [8,12]. This is an expected finding because the detection of dark and light targets is inferred to be via the OFF pathway and ON pathway respectively. The OFF pathway has a higher contrast gain than the ON pathway as shown in visually evoked potential (VEP) recordings in humans [13]. Besides, in animal models such as cats, it has been identified that OFF neurons have a larger representation in the visual cortex [3] and stronger connections in the thalamus relative to the ON neurons [4]. Also, the OFF pathway consists of almost twice the number of cortical neurons compared to the ON pathway [5]. Recently, the ‘neuronal blur’ hypothesis has been proposed to explain why it is harder to detect light targets than dark targets using a centre-surround model [14]. It states that the enlargement of the image due to ON response luminance saturation [7] causes the image to fall in the surround region, thus enhancing the suppression [14]. The ON/OFF asymmetries are also noticed at the retinal level [6,15–18].

This is the first study that has looked at visual performance in the detection of light targets and dark targets over a wide age range of subjects from 18 years to 80 years. There was a significant correlation of reaction time with age across the entire age range. However, > 49 years of age, the reaction time for both targets (dark and light) were not correlated with age. Both these findings are consistent with Zhao et al (2014) [8]. Across all ages and in the subgroups (≤ 49 years and > 49 years), we have found that accuracy for both light and dark targets are independent of age, which is similar to the previous study [8].

Dark targets on light background stimulate the OFF pathway whereas, the light targets on dark background stimulate the ON visual pathway [14]. Westheimer et al (2003) had measured acuity on the normal polarity and reverse polarity charts on healthy normal subjects as a function of age [19], which is essentially a test of ON and OFF visual pathways. Acuities measured on both the charts showed a negative correlation with age. They showed that the rate of change in acuity measured using normal polarity charts (OFF pathway) is steeper than the acuities measured on the reverse contrast chart [19]. However, the rate of deterioration in visual performance for dark targets (OFF pathways) was similar to the light targets (slopes in Fig 3E and 3F) in this study. The discrepancies could be attributed to the differences in tasks (detection task [this study] Vs recognition task [Westheimer, 2003] measured between the studies. Besides, it is well known that reaction time in search-based tasks, increases with ageing [20]. The role of eye-hand coordination in increased reaction time with age cannot be discounted [21]. However, it could also be attributed to neuronal loss in the visual cortex [22]. There is currently no evidence to suggest that ageing affects one pathway more than the other in humans. None of our patients had any motor or neurological problems that could influence eye-hand coordination.

The differences between reaction time for light and dark targets showed no correlation with age, which is contrary to the previous report. Similarly, asymmetry in reaction times between ON and OFF targets (∼200 ms) was also not very pronounced compared to Zhao et al [8] (∼500 – 600 ms). The average time taken for identifying the dark targets is greater than the dark targets reaction time found in Zhao et al [8]. However, the magnitude of difference in accuracy between light and dark targets was smaller (∼ 2 - 3 units) compared to the previous study reported by Zhao et al [8] which was ∼8 %. The major difference between this study and the lab-based study is that they had performed on a desktop monitor, while this was on an iPad. Therefore, we replicated the test containing 1 - 3 dark or light targets in a randomized manner using Psychopy program [23] and the participant was asked to press the keyboard buttons to respond. The results from this small sub-sample (n=10) were similar to the larger database. The asymmetry in reaction time and accuracy was still smaller between the light and the dark targets. Although we tried to replicate the setup done by Zhao et al [8], there could be some methodological differences that could have accounted for the discrepancies. It is known that at higher luminances, the asymmetry in reaction times and accuracy between light and dark targets reduces [14]. That could also be attributed to the different results.

The normative database forms a useful test to differentiate age-related losses from disease-specific losses. The pathway-specific losses are noticed in glaucoma [8], melanoma-associated retinopathy [9], congenital stationary night blindness and also expected in rod-dominated diseases such as retinitis pigmentosa, rod dystrophy and rod monochromatism. The rationale for expected loss in rod-dominated diseases is based on the previous findings that ON pathway defects are known to be associated with rod system dysfunction [24]. Therefore, the ON pathway function can potentially provide a surrogate marker of rod function, which is typically difficult to measure without dark adaptation and elaborate individual calibration [25].Thus the implications of the test go well beyond the assessment of ON and OFF visual pathways.

The limitations of the study are that the test was not carried out in a controlled lab-based setup. The external lighting under which, the test was conducted was not uniform across all the participants. However, such effect on the results is expected to be minimal, as the test is primarily dependent on the luminance of the iPad screen. The angle at which the device held during the test was not fixed. However, most participants would hold at an angle that they would hold reading material. Also, there is no fixation monitor to track participants’ eye movements, however, the subjects were instructed to keep looking towards the centre of the screen throughout the test. There was no ‘pause’ option available during the test. Therefore, the effect of fatigue on reaction times cannot be ruled out.

## Conclusions

The test has several advantages such as easy-to-use, portability and being less time-consuming. Therefore, having an age-related normative database along with it would make it more attractive for a wider usage to distinguish age-related changes from disease-specific defects. Future studies should iterate and identify the optimum ‘number of trials’ required to balance between the need for robust data and the time taken for test completion. The next step should be testing in retinal diseases, to see if the indices can provide additional diagnostic information, such as ON pathway deficits in rod-dominated diseases.

## Acknowledgements

We would like to acknowledge Hyderabad Eye Research Foundation for the infrastructure support to carry out the project.

